# VcfR: a package to manipulate and visualize VCF format data in R

**DOI:** 10.1101/041277

**Authors:** Brian J. Knaus, Niklaus J. Grünwald

**Affiliations:** Horticultural Crops Research Unit, USDA-ARS, Corvallis, 97330, USA; Department of Botany and Plant Pathology, Oregon State University, Corvallis, OR, 97331, USA

**Keywords:** data visualization, high throughput sequencing, quality control, VCF format

## Abstract

Software to call single nucleotide polymorphisms or related genetic variants has converged on the variant call format (VCF) as the output format of choice. This has created a need for tools to work with VCF files. While an increasing number of software exists to read VCF data, many only extract the genotypes without including the data associated with each genotype that describes its quality. We created the R package vcfR to address this issue. We developed a VCF file exploration tool implemented in the R language because R provides an interactive experience and an environment that is commonly used for genetic data analysis. Functions to read and write VCF files into R as well as functions to extract portions of the data and to plot summary statistics of the data are implemented. VcfR further provides the ability to visualize how various parameterizations of the data affect the results. Additional tools are included to integrate sequence (FASTA) and annotation data (GFF) for visualization of genomic regions such as chromosomes. Conversion functions translate data from the vcfR data structure to formats used by other R genetics packages. Computationally intensive functions are implemented in C++ to improve performance. Use of these tools is intended to facilitate VCF data exploration, including intuitive methods for data quality control and easy export to other R packages for further analysis. VcfR thus provides essential, novel tools currently not available in R.

## Introduction

Genetic sequence variations are stored in variant call format (VCF) data files for bioinformatic and genetic analysis. Bioinformatic tools for calling variants such as SAMtools [16] or the GATK’s haplotype caller [1, 6, 18] have all converged on VCF [5, 23] as an output file format. VCF files have become the de facto standard for downstream analyses of variant data. Prior to analysis, genetic variants should be screened and filtered based on quality metrics provided by the variant callers such as SAMtools or GATK-HC. This is particularly important for non-model organisms where curated panels of variants are not currently available. Convenient and interactive tools for manipulating these data are not currently available in the R language, a popular environment for working with genetic data.

A VCF file consists of a text file structured according to the VCF definition [5, 23]. These files may be provided as text files or as compressed files, most typically in the gzip format. Each file begins with a meta region where abbreviations used in the data portion of the file are defined with one definition provided per line. The tabular data portion of the file includes samples in columns and variants in rows. The first eight columns contain information that describes each variant. The ninth column contains the FORMAT specification for genotype records. All subsequent columns contain sample information where each column corresponds to a sample and the values within each column consist of genotypes and any optional information specified by the FORMAT column. This definition allows a flexible amount of information to be included with each genotype. Variant data can include a SNP, deletion, insertion, large structural variants and/or complex events. However, because the data are not strictly tabular, it presents a challenge in that it needs to be parsed. The quantity of data contained in these files requires this parsing to be performed in an efficient and reproducible manner.

Among currently existing tools for working with VCF files is a collection of tools called VCFtools [5]. VCFtools provides an extensive set of tools for data filtering and analysis. Because they are command line tools they are ideal for high performance computing environments which lack graphical user environments or are implemented in cloud-based queueing systems lacking interactive visualization. However, some users may prefer a more interactive set of tools. Frequently, a set of quality control criteria are used to filter data with no validation of how these criteria may affect the resulting dataset. A graphical and interactive method to manipulate these files would allow researchers to rapidly determine how choices in filtering may affect the resulting dataset.

We present vcfR, an R package designed to help users manipulate and visualize VCF data. Functions are included that efficiently read VCF data into memory and write it back to disk. A parsing function efficiently extracts matrices of genotypes or their associated information. Plotting functions provide a rapid way to visually assess variant characteristics. Because this software is implemented in R it also provides ready access to the multitude of statistical and graphical tools provided by the R environment. Through efficient parsing and visualization, vcfR provides a tool to rapidly develop hard filters for quality metrics that can be easily tailored to individual projects and experimental designs. Key components of vcfR are implemented in C++ and called from R to minimize time of computation. VcfR is open source and available on CRAN and GitHub with appropriate documentation.

## Materials and Methods

### The pinfsc50 example data set

An example data set for the diploid, oomycete pathogen Phytophthora infestans is provided to demonstrate application of vcfR. We use sequence data from supercontig_1.50 in the FASTA format from the published genome [11, 2] as example sequence data. We also provide an annotation file in the GFF format for supercontig_1.50. Lastly, we provide the variant data in VCF format. VCF format data is based on short read data from previously published sources [4, 2, 17, 29]. This short read data was mapped to the T30-4 reference with bwa-mem [15], while bam improvement and variant calling was performed according to the GATK’s best practices [1, 6, 18]. Phasing was performed with beagle4 [3]. Because beagle4 removes most of the diagnostic information output by the GATK’s haplotype caller, the phased genotype from the beagle4 VCF file was appended to the haplotype caller’s VCF file (after the unphased genotype was removed), resulting in the VCF file provided. It is important to note that we have processed our own data as many publicly available datasets lack the richness of descriptors provided by variant calling software. For example, data available from the 1,000 genomes project only contains the genotypes and none of the quality metrics (i.e., it is a production dataset). By providing our own processing it allows for provision of VCF format files rich with information that can be used to provide instructive examples.

Ideally, this example dataset will be somewhat large so that it demonstrates efficient execution of functions intended to be used on typically larger VCF data sets. However, packages hosted on CRAN currently generate a ‘NOTE’ when package size exceeds 5MB. This arbitrary threshold creates a practical limitation to ensure a data set is small enough to distribute efficiently. In order to balance this need for size we have released the pinfsc50 dataset on CRAN as its own R package with the same name and hope others will find it useful as well.

### Efficient file access

A common bottleneck to data intense projects is reading and writing files from disk to memory, and back to disk. Essential criteria for VCF file input and output include that it must be fast, able to read text and gzipped files, and able to handle the non-tabular VCF format. Furthermore, input ideally should allow subsetting to specific rows (variants) and columns (samples) of interest. This allows work to be performed on datasets where the entire file may require more physical memory than is available in a given computational environment. We created a custom VCF file reading function using Rcpp [9] to execute the computationally intensive steps in C++. This C++ code allows efficient reading and writing of VCF files.

To provide a comparison among our new function and existing R functions we conducted a benchmarking test. Evaluation was run on a 3.60GHz Intel® Core™ i7-4790 CPU running Ubuntu 12.04.5 LTS with a Western Digital® WDC WD10EZEX-75M drive. Our test data set, pinfsc50, was a gzipped VCF file containing 29 meta lines, 22,031 variants and 27 columns (18 samples). Functions to read tabular data (*utils∷read.table*() and *data.table∷fread*()) were parameterized to skip the non-tabular meta region, providing a slight advantage to these functions. The function *data.table∷fread*() was called as *data.table∷fread*(*’zcat filename.gz’*) because it does not currently read in gzipped data. Data input was run twice to determine if functions implemented some form of caching that may improve a second read time. Input time was measured with the R function *system.time*() where elapsed time was recorded.

### Manipulation of VCF data

Once the VCF data is read into memory, manipulation of the data follows. The data for each genotype from a VCF file can be seen as a colon delimited string containing the genotype as well as optional information to characterize each genotype. The FORMAT column specifies the format of these elements. For example, many variant callers will provide information on how many times each allele was sequenced, genotype likelihoods and other information to characterize the quality of each genotype. An example of a FORMAT element and a genotype element is provided in equation 1:

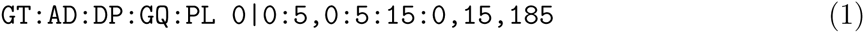

Here ‘GT’ is the genotype (0|0, homozygous for the reference) and allelic depth (AD) for the genotype is ‘5,0’ (the reference allele was sequenced five times and the alternate allele was sequenced zero times). The definition of these abbreviations should be found in the meta region of the file. As the number of variants and samples increases, the task of parsing this data increases multiplicatively. This presents a performance challenge to typical R functions. We included a parser (the function *extract.gt*()) that is implemented using Rcpp [9] to rapidly extract this information. The result is a matrix of strings or numerics corresponding to one of the specified FORMAT values. These matrices can then be manipulated as typical R matrices and be visualized using the base R graphics system [21] or other tools.

### Chromosomal summaries

Variants for VCF data can be incorporated with sequence and annotation data to provide chromosomal perspectives. Within the vcfR framework this process results in the creation of an object of class chromR. The vcfR function *chromoqc()* can be used to visualize chromR objects. During the creation and subsequent processing of this object, summary statistics of the variants are provided (heterozygosity, effective size) and sliding window analyses are performed (nucleotide content, number of annotated positions, variants per window). These per variant and per window summaries are stored in a tabular format that can be visualized with the R base graphics system (e.g., *graphics∷hist()* and *graphics∷plot()*) or saved to a file. A genomic perspective can then be obtained by concatenating chromosomal summaries. Integration of variant data with sequence and annotation data provides a novel tool to rapidly identify genomic regions of interest.

## Results

The vcfR package includes novel functions for reading and writing data from VCF files and for visualizing, manipulating and quality filtering of data.

### File access

We wrote code for reading and subsetting VCF data in C++ to improve computational speed. Results from the file access benchmarking test are presented in table 1. In general, the typical R functions for data input, such as *utils∷read.table()* do not perform well. The function *utils∷read.table()* required 2-3 seconds to read in our modest size data set (18 samples, 22 thousand variants). More efficient methods of reading tabular data into R include *read_table()* in the package readr [28] and *fread()* in the package data.table [8]. The function *data.table∷fread()* is perhaps the best performing function with the greatest flexibility in terms of column and row access. It read data into memory in 0.1-0.2 seconds. However, it lacks the ability to read gzipped files and is designed for reading only tabular data. On Unix systems, gzipped data may be read in by modifying the call to include *zcat* or *gzcat*. Windows users may be required to make an uncompressed version of their file to facilitate use, resulting in redundancy of files, a practice that is unnecessary. These general functions that read in tabular data may provide a path for data input, however, accessing the resulting data for manipulation from the resulting table remains an issue. Because each element read in by these functions that read in tabular data may contain data associated with each genotype as a colon delimited string they need further processing before attaining a format for analysis in existing R genetics packages.

**Table 1:** Comparison of R functions that read VCF or VCF-like data into R. The pinfsc50 data used for performance benchmarking consisted of 18 samples and 22,031 variants in a gzipped VCF file.

|  | **utils::**<br>**read.table()** | **readr::**<br>**read_table()** | **data.table::**<br>**fread()** | **pegas::**<br>**read.vcf()** | **VariantAnnotation::**<br>**readVcf()** | **vcfR::**<br>**read.vcfR()** |
| --- | --- | --- | --- | --- | --- | --- |
| Reads VCF | F | F | F | T | T | T |
| Reads gzip | T | T | F | T | T | T |
| Conversion functions | F | F | F | T | F | T |
| Row selection | T | T | T | T | T | T |
| Column selection | F | F | T | F | T | T |
| First read (sec) | 3.259 | 0.637 | 0.257 | 2.260 | 5.739 | 0.419 |
| Second read (sec) | 2.413 | 0.384 | 0.177 | 0.387 | 1.246 | 0.377 |

In addition to the functions already mentioned above, functions currently exist to read VCF files into R. The package PopGenome includes the function *readVCF()*. This function reads in VCF files that are both faidx indexed and bgzipped, making this function beyond the scope of these comparisons. This may present a direction high throughput projects may take in the future. At the present, it is our experience that gzipped files are more common. This package also does not appear to include conversion functions to translate data into data structures used in commonly used R packages such as ape [20], adegenet [12], pegas [19] and poppr [14, 13]. This function therefore appears applicable to only this package. The Bioconductor package VariantAnnotation includes a *readVcf()* function. It does not appear to perform as well as the other options at reading in data as it required 5.7 and 1.2 seconds to read in data (Table 1). Also, the resulting data structure appears complicated relative to the other options presented here and appears specific to the Bioconductor project. Users who are already invested in the Bioconductor data structures may find this to be a viable route. If the user is instead interested in analyses from packages from outside of Bioconductor (e.g., CRAN), this route is less appealing. The package pegas includes the function *read.vcf()*. This function appears to fulfill all of our criteria. However, this function only reads in the genotypes from the VCF file and none of the data associated with each genotype. This may be because the object of class loci, created by this function, does not support associated information. This means that this function will not provide information for quality filtering, a task crucial to obtaining high quality data. This function read in our test file in 2.3 and 0.4 seconds. The first read is comparable to *utils∷read.table*(), suggesting relatively poor performance.

However, the second read was comparable to *data.table∷fread* and *vcfR∷read.vcfR*, suggesting a well performing input function. The difference in input speed suggests that some sort of caching may be occurring that makes subsequent reads much faster than initial reads for this function. Our function *vcfR∷read.vcfR()* performs comparably to *data.table∷fread()* and *readr∷read_table()* (Table 1) but provides the convenience that it is specifically designed to handle VCF data whether compressed (gzip) or not. It also provides the ability to select rows and columns from a VCF file so that partial files can be read in to conserve memory. Lastly, this package provides conversion functions to translate data from the vcfR object created by *vcfR∷read.vcfR()* into formats supported by other R genetics packages. Most importantly, as described below in more detail, our VCF handling includes quality metrics available in input files for subsequent filtering.

### Data parsing and visualization

Once VCF data are read into memory they are ready for further processing. However, because the data are not in a strictly tabular format (see equation 1) further processing is needed to access the data. Our function *extract.gt()* parses elements from the gt portion of VCF data. For example, the read depth at each variant (DP) is commonly included in VCF files either by default or as an option. Invocation of *extract.gt()* on VCF data containing depths results in a numeric matrix of depths with samples in columns and variants in rows. This data can then be visualized with existing R tools, such as ggplot2 [27] which was used to summarize a matrix of depths using violin plots (Figure 1). Similarly, a modified heatmap function provided in vcfR (*heatmap.bp()*) can be used to summarize the depth matrix (Figure 2). The steps to achieve these results are illustrated with the code provided below.

**Figure 1:**
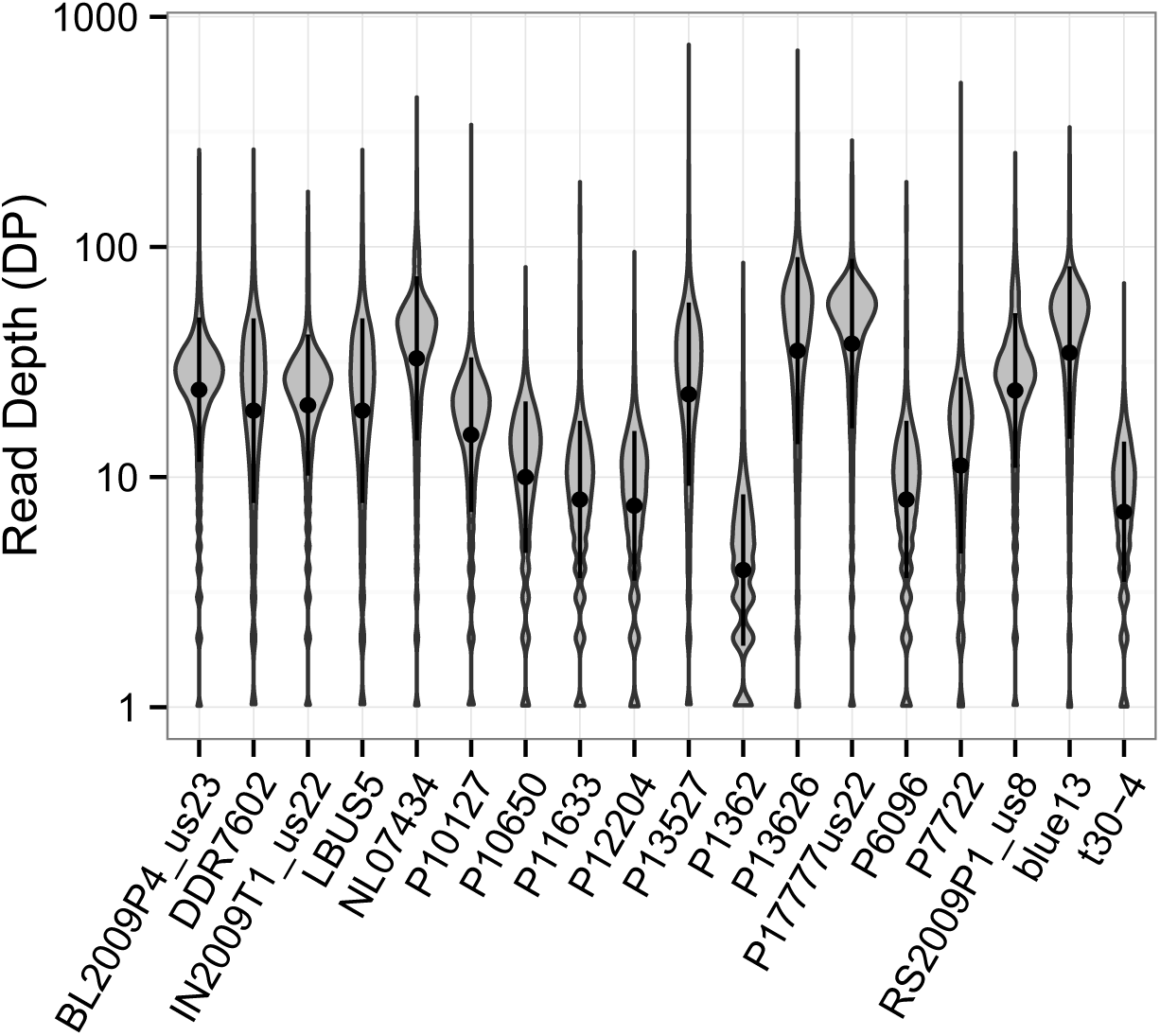
Violin plot of read depth (DP) for the 18 samples in the pinfsc50 data set. A numeric matrix was produced from the VCF file with the function *vcfR∷extract.gt()*.

**Figure 2:**
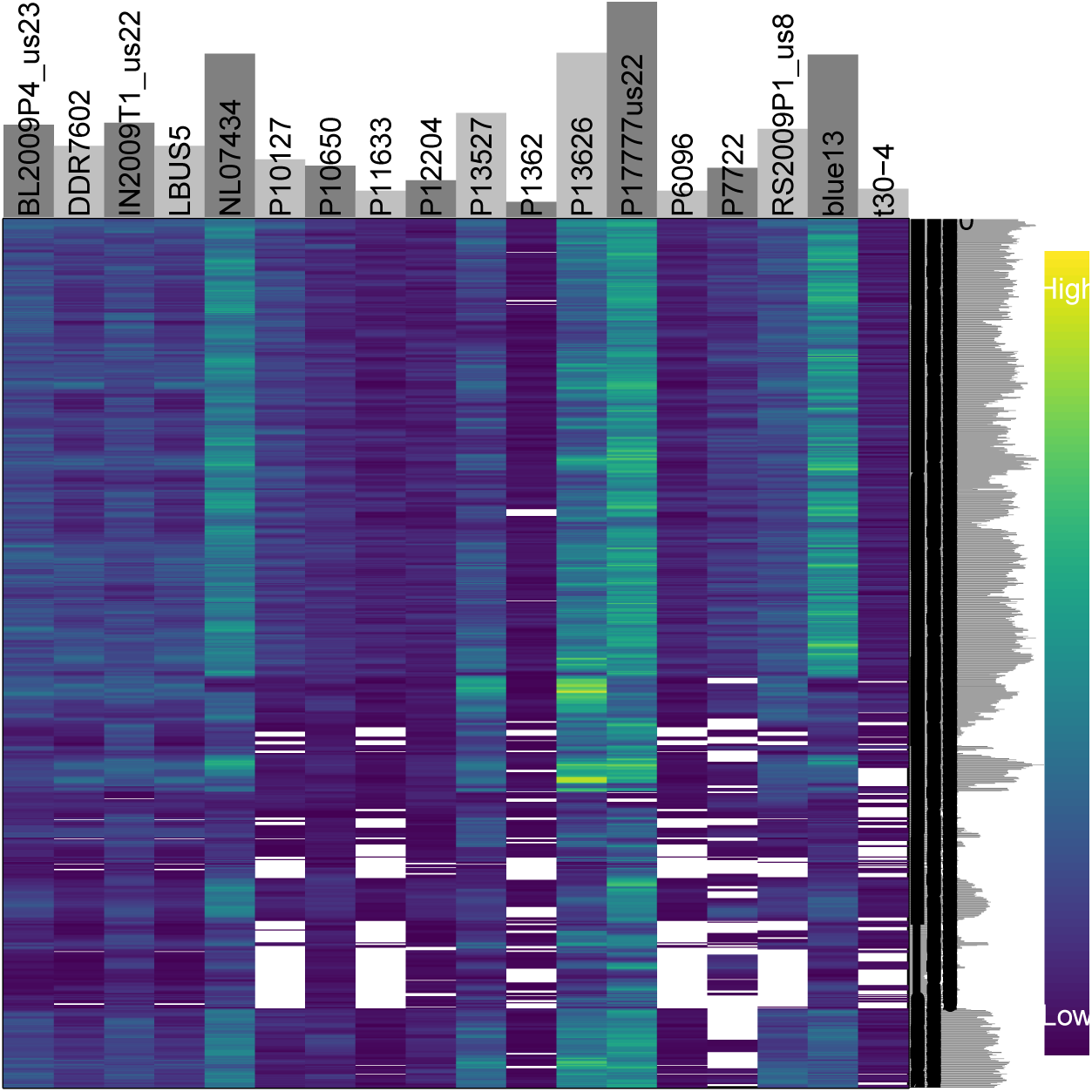
Heatmap of read depth (DP) for the 18 samples in the pinfsc50 data set. Each column is a sample and each row is a variant. The color of each cell corresponds to each variant’s read depth (DP). Cells in white contain missing data. Marginal barplots summarize row and column sums. The matrix was extracted from the VCF data with the function *vcfR∷extract.gt()* and the plot generated with *vcfR∷heatmap.bp()*.

~~~
# Load libraries.
library(vcfR)
library(pinfsc50)
library(reshape2)
library(ggplot2)
# Find and input VCF data.
vcf <- system.file("extdata", “pinf_sc50.vcf.gz", package = “pinfsc50")
vcf <- vcfR∷read.vcfR(vcf, verbose = FALSE)
# Parse DP from the gt region.
dp <- extract.gt(vcf, element="DP", as.numeric = TRUE)
# Reorganize and render violin plots.
dpf <- melt(dp, varnames=c("Index", “Sample"), value.name = “Depth", na.rm=TRUE)
dpf <- dpf [dpf$Depth > 0,]
p <- ggplot(dpf, aes(x=Sample, y=Depth)) + geom_violin(fill="#C0C0C0", adjust=1.0,
                                                                       scale = “count", trim=TRUE)
p <- p + theme_bw()
p <- p + ylab("Read Depth (DP)")
p <- p + theme(axis.title.x = element_blank(),
                             axis.text.x = element_text(angle = 60, hjust = 1))
p <- p + stat_summary(fun.data=mean_sdl, geom="pointrange", color="black") p <- p + scale_y_continuous(trans=scales∷log2_trans(), breaks=c(1, 10, 100, 1000))
p
# Plot as heatmap.
heatmap.bp(dp[501:1500,])
~~~

This functionality adds data parsing to file input in vcfR thereby creating an easy path to visualization of VCF data.

### Quality filtering

Variant calling software typically requires some form of post-hoc quality filtering. This is particularly important in non-model systems that contain only a small portion of curated data [10]. The vcfR function *chromoqc()* was used to generate Figure 3 from a FASTA sequence file, a GFF annotation file and a VCF variant file. Inspection of Figure 3 illustrates a number of issues typically observed in raw VCF data. For example, the read depth (DP) is highly variable. The marginal box and whisker plot for the read depth panel contains 50% of the data within the boxes of the box and whisker plot. This provides an estimate of depth to expect for base ploid variants. Below this region may be variants of low coverage that may not have been called accurately. Above this region are variants that may be from repetitive regions of the genome and may therefore violate ploidy assumptions made by the variant caller. Mapping quality (MQ) consists primarily of variants with values of 60 as well as a population of variants of a lower quality. By using the vcfR function *masker()* we can filter on thresholds for these values. Here we have used a read depth between 350 and 650 and a mapping quality between 59.5 and 60.5. The result is visualized in Figure 4. We see that we eliminated most of the variants on the right hand side of the plot. The resulting data set may now be considered to be of higher stringency. Alternatively, a researcher may want to use this variant set to use as a training set for another round of variant discovery. Processing and visualization of these results can occur with a few lines of code:

**Figure 3:**
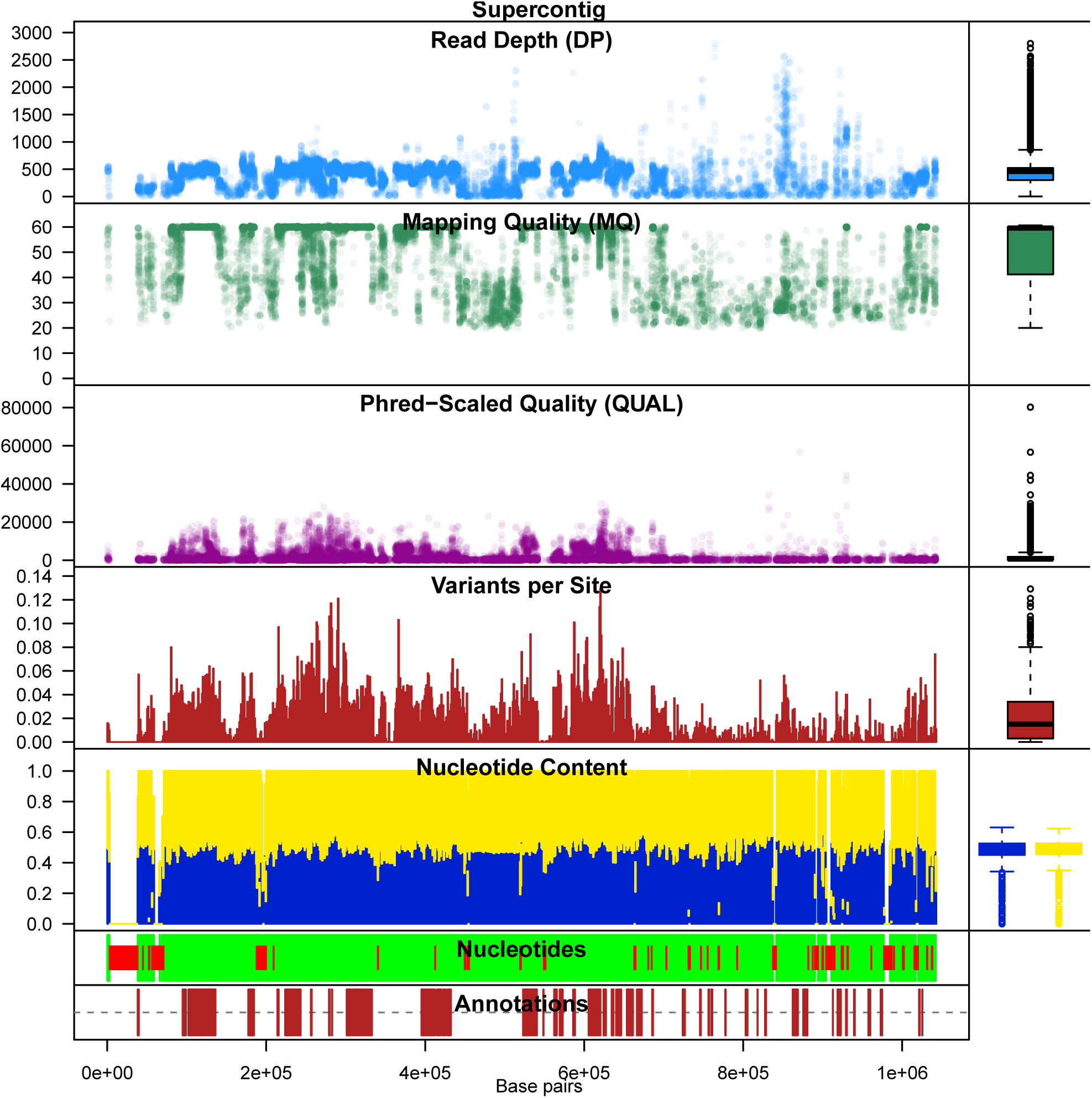
Chromoqc plot showing raw VCF data for one supercontig in the pinfsc50 data set. The lowest panel represents annotations as red rectangles. Above this is a panel where regions of called nucleotides (A, C, G or T) are represented as green rectangles and ambiguous nucleotides (N) are represented as red, narrower, rectangles. Continuing up the plot is a sliding window analyses of GC content and then one of variant incidence. Above these are three dot plots of phred-scaled quality ), mapping quality (MQ) and read depth (DP).

**Figure 4:**
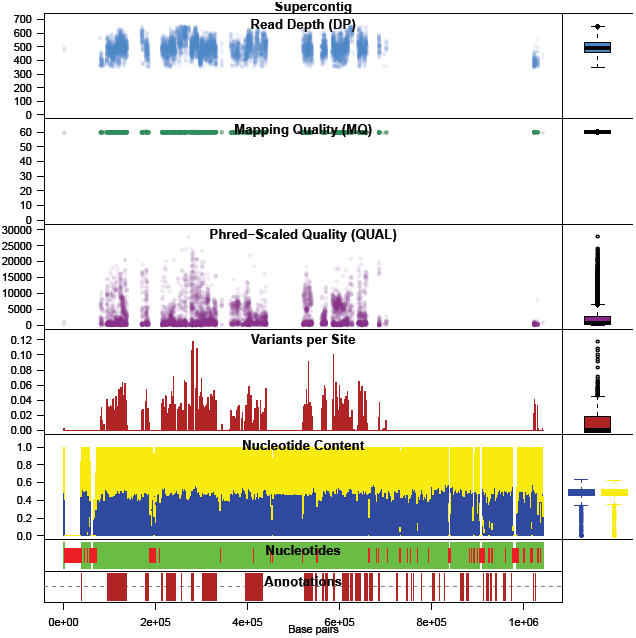
Chromoqc plot showing the one supercontig in the pinfsc50 data set after filtering on read depth (DP) and mapping quality (MQ).

~~~
# Load libraries
library(vcfR)
library(pinfsc50)
# Determine file locations
vcf_file <- system.file("extdata", “pinf_sc50.vcf.gz",
                               package = “pinfsc50")
dna_file <- system.file("extdata", “pinf_sc50.fasta",
                               package = “pinfsc50")
gff_file <- system.file("extdata", “pinf_sc50.gff",
                               package = “pinfsc50")
# Read data into memory
vcf <- read.vcfR(vcf_file)
dna <- ape∷read.dna(dna_file, format = “fasta")
gff <- read.table(gff_file, sep="\t", quote="")
# Create a chromR plot
chrom <- create.chromR(name="Supercontig",
                              vcf=vcf, seq=dna,
                              ann=gff)
# Mask for read depth and mapping quality chrom <- masker(chrom, min_QUAL=0, min_DP=350,
                              max_DP=650, min_MQ=59.5,
                              max_MQ=60.5)
chrom <- proc.chromR(chrom)
# Plot.
chromoqc(chrom, dp.alpha=20)
~~~

### Data export

Data can be exported from vcfR into several data formats useful for downstream analysis. The most straight forward option may be to output the manipulated data as a VCF file. The function *write.vcf()* takes a vcfR object and writes it to file as a gzipped VCF file. This allows any software that operates on VCF files to be used for downstream analysis. If the researcher prefers to remain in the R environment, several other options exist. The function *vcfR2genind()* can be used to convert modest amounts of VCF data into a genind object, allowing analysis in adegenet [12]. Genind objects may be easily converted to genclone objects with *poppr∷as.genclone()* for analysis in poppr [14, 13]. Authors of the adegenet package have more recently created the genlight object specifically for high throughput sequencing applications. Objects of class genlight can be created from objects of class vcfR using the function *vcfR2genlight()*. Once an object of class genlight has been created it can be converted to an object of class snpclone using *poppr∷as.snpclone()*. When sequence information is provided, the VCF data can be converted into an object of class DNAbin using *vcfR2DNAbin()* for analysis in ape [20] or pegas [19]. The inclusion of data conversion functions allows VCF data to be easily converted into data structures used by currently existing R genetics packages making these existing methodologies available to the analysis of VCF data.

## Discussion

The advent of high throughput sequencing has provided researchers with a deluge of data. As with all data, some of it is of high quality while some of it may not be. The VCF file format provides a flexible format that authors of variant callers can use to include a diversity of information to support genotype calls. However, software available to utilize this information, particularly in the R environment, is currently limited.

The R package vcfR is a novel tool for manipulation and visualization of data contained in VCF files. This package contains functions to efficiently read and write VCF data from and to files. Functionality is also provided to parse VCF data once loaded into memory. This creates an entry point for VCF data analysis in the R environment with its associated genetic analysis packages. For example, functions in vcfR can read in VCF data, extract numeric values such as read depth or genotype qualities from this data, and the data can then be visualized using standard R scatter plots or histograms. These data can also be visualized with custom plots provided by vcfR. This information can be used to determine thresholds for quality filtering, similar to VCFtools [5], but in a graphical, interactive R environment. The package also includes functions to convert this information to formats used by existing R packages specifically designed to work with population genetic data (e.g., ape [20], adegenet [12], pegas [19] and poppr [14, 13]). Once VCF data is read into memory, typically a single function is all that is required to translate the data into a data structure supported by these other packages. This makes the data contained in VCF files available to functions provided by the vcfR package, R’s standard plotting functions as well as methodologies that currently exist in available R packages.

The integration of vcfR with existing methodologies provides researchers with a rapid path towards research products. Once VCF data have been generated, they can be rapidly queried for quality metrics to determine quality filtering thresholds. This information can then be used to filter data within vcfR or the thresholds can be used in server side processes such as VCFtools [5]. Once VCF data has been determined to be of sufficient quality for downstream use, vcfR provides data export tools for other analytical tools available in R. This novel functionality allows VCF data to be easily handled from data acquisition, through quality control and final analysis within the R programming environment. VcfR provides flexibility to aid researchers explore quality thresholds and other analytical decisions in an efficient manner to ensure the lowest amount of technical variation and the highest quality results.

## Acknowledgements

We acknowledge Javier F. Tabima (Department of Botany and Plant Pathology, Oregon State University) for extensive alpha testing that greatly improved the R code. Zhian N. Kamvar (Department of Botany and Plant Pathology, Oregon State University) provided valuable discussion on R package development. N.J.G. thanks NESCent for organizing the 2015 Population Genetics in R Hackathon in Durham, NC. This research is supported in part by US Department of Agriculture (USDA) Agricultural Research Service Grant 5358-22000-039-00D and USDA National Institute of Food and Agriculture Grant 2011-68004-30154 to N.J.G.

## Data Accessibility

The program and user manual are available on CRAN (cran.r-project.org/package=vcfR). The pinfsc50 dataset is available on CRAN (cran.r-project.org/package=pinfsc50).

## Author Contributions

BJK conceived of the project, wrote code, wrote the documentation, and wrote the manuscript. NJK coordinated the collaborative effort, discussed interpretation, wrote the manuscript and obtained funding.

